# Decoding nonspecific interactions between human nuclear transport proteins: A computational study

**DOI:** 10.1101/2021.03.22.436462

**Authors:** Shravan B. Rathod

## Abstract

The nuclear protein transport between the nucleus and cytosol can be considered a core process of cell regulation. Specially designed proteins in nature such as importins, exportins, and some other transporters facilitate this transport in the cell and control the cellular processes. Transient and weak protein–protein interactions are basis of these various biomolecular processes. Prior to cargo transports, the transport proteins recognize the Nuclear localization signals (NLSs) and Nuclear export signals (NESs) of cargo proteins and, bind to the RanGTP. Also, these proteins bind with other similar protein subunits along with RanGTP to transport cargos. Cell is enormously crowded place where DNA, RNA, proteins, lipids and small molecules cooperatively facilitate numerous cellular processes. In such environment, existence of nonspecific interactions between proteins is quite obvious. Considering this hypothesis, in this study, protein-protein docking approach was applied to determine the binding affinities of 12 human nuclear transport proteins. Results showed that KPNA1, TNPO1 and TNPO3 have greater affinity to bind with other transport proteins. Also, among 78 complexes (12 homodimers and 66 heterodimers), KPNA1-KPNB1, KPNA1-TNPO1 and KPNA1-TNPO3 complexes have the highest stability.

**Graphical abstract:** 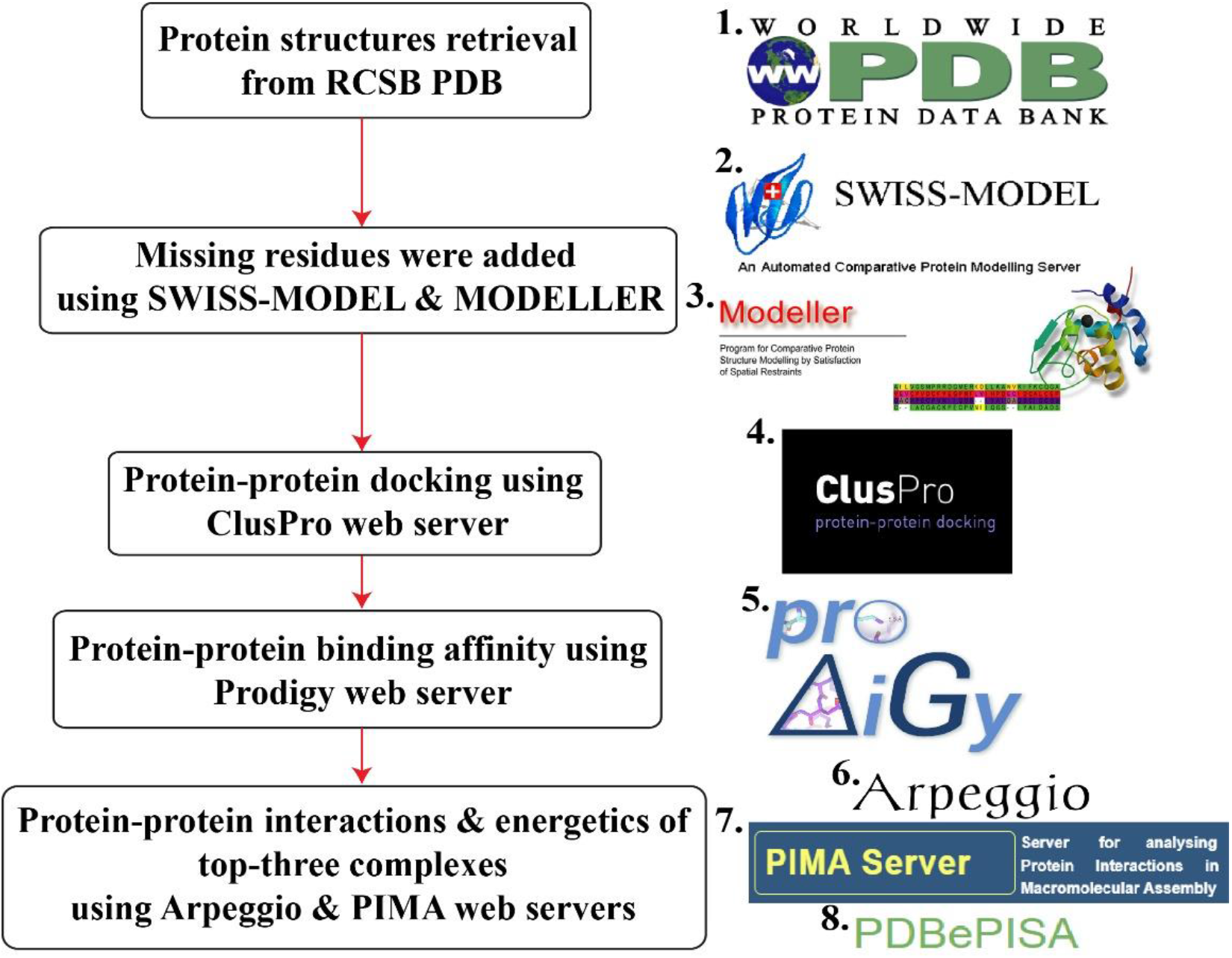

Initially, 12 human nuclear transport proteins PDB structures were retrieved from the 1. Protein data bank (PDB). These proteins had some missing terminals and residues thus, we used 2. SWISS-MODEL and 3. MODELLER v.10.1 to model those regions in these proteins. Next, we used widely popular web server, 4. ClusPro v.2.0 for protein-protein docking analysis among 12 proteins. Then, we employed 5. PRODIGY web server to calculate the binding affinities of 78 complexes (12 homodimers & 66 heterodimers). Finally, we utilised three web tools, 6. Arpeggio, 7. PIMA and 8. PDBePISA to analyse top-three complexes (KPNA1-KPNB1, KPNA1-TNPO1 & TNPO3) for in-depth interactions and energetics.

## Introduction

Cargo transportation between the nucleus and cytoplasm is an essential process for the cellular regulation and plays critical role in diverse physiological mechanisms such as cellular signal transduction, gene expression, and cell differentiation [1-2]. In eukaryotic cells, nuclear and cytoplasmic contents are physically separated by the lipid bilayer of nuclear envelope. Envelope has a large number of Nuclear pore complexes (NPCs) through which many molecular components make their journey between the nucleoplasm and cytoplasm. NPCs are composed of the assembly of proteins called nucleoporins [3-4]. Large and polar proteins cannot be transported via the diffusion process hence some selective transporters help them to commute between the cytoplasm and nucleus through NPCs [5-6].

Transporters (importins and exportins) recognize the specific sequence to bind in cargo proteins called Nuclear localization signal (NLS) or Nuclear export signal (NES) and form a complex to transport them at a certain spot in the nucleus or cytoplasm. But, this is not the full story because to drive these processes, another important protein called RanGTP (Ras-related nuclear protein guanosine-5’-triphosphate) binds to the transporter and supplies energy to transport cargos [7]. Hence, the complex formation between the cargos, transporters, and RanGTP is an inevitable step in protein transports.

Research revealed that a single eukaryotic cell has approximately 42 million proteins thus, cell is considered extremely crowded place [8]. Owing to these huge number of proteins, it can be thought that many proteins can bind to other proteins non-specifically. These nonspecific interactions between the proteins trigger numerous diseases-linked pathways [9-10]. Hence, it is possible that in the cytoplasmic and nucleoplasmic molecular crowding, transporters can have nonspecific binding efficiency with each other and to form a homo-or heterodimer. To determine the nonspecific interactions between the proteins, techniques such as NMR-based Paramagnetic relaxation enhancements (PRE) [11] and nanoflow electrospray ionization mass spectrometry (nanoES-MS) [12] have been developed. In this present study for the primary investigation of nonspecific bindings between human nuclear transport proteins, computational analyses were performed. In the beginning, molecular docking between the 12 human nuclear transport proteins was carried out using protein-protein docking algorithm. Then, to determine the binding energies and interactions of complexes, multiple web tools have been employed. Additionally, Electrostatic potential (ESP) surface analysis of each protein was done to get further insight into the protein-protein bindings and interface interactions.

### Materials and methods Protein preparation

The crystal structures of 12 proteins were retrieved from the Protein data bank (PDB) (https://www.rcsb.org/). Only human nuclear transport proteins were selected and preferred on the basis of their high-resolution structures with zero mutations. Heteroatoms such as solvent molecules and ligands were removed during protein preparation in PyMOL v.2.5.2 [13]. Further, the missing regions of these 12 proteins were modelled using SWISS-MODEL [14] available at https://swissmodel.expasy.org/ and MODELLER v.10.1 [15]. Next, the missing hydrogens in protein structures were added using PyMOL. PDB code, sequence length, and crystallographic resolution of proteins are shown in Table 1 whereas Fig. 1 illustrates the structures of 12 human nuclear transport proteins.

**Table 1.**
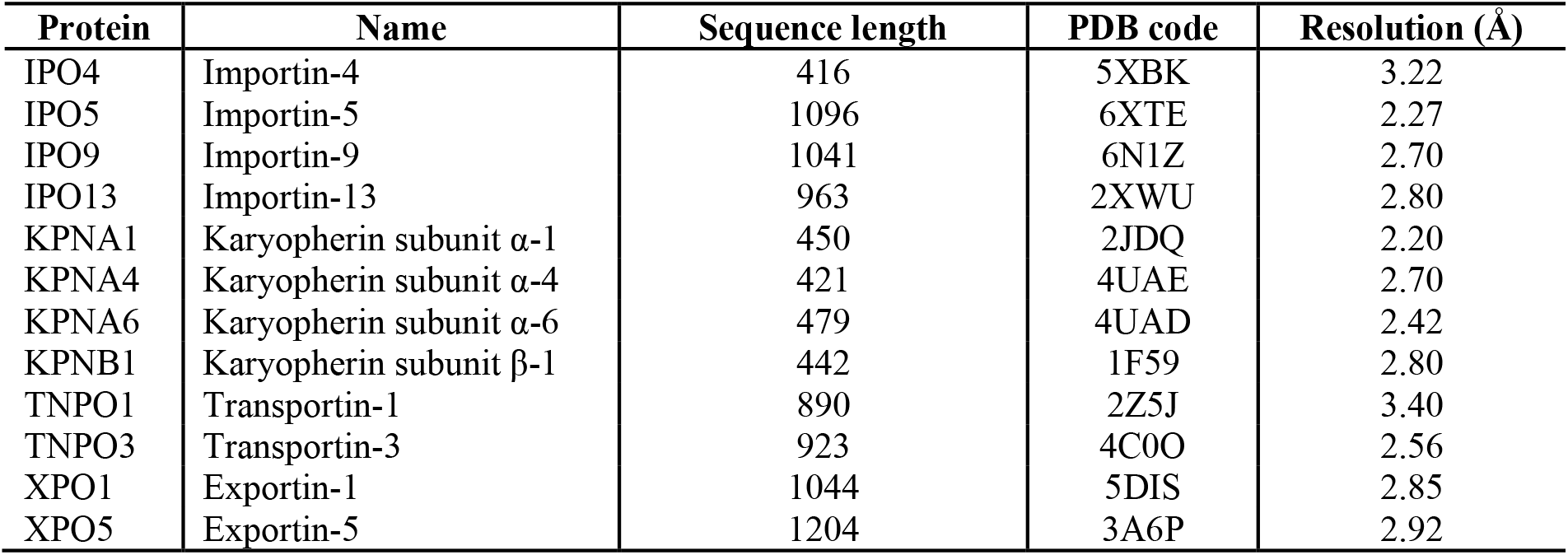
Structural information of 12 human nuclear transport proteins.

| Protein | Name | Sequence length | PDB code | Resolution (Å) |
| --- | --- | --- | --- | --- |
| IPO4 | Importin-4 | 416 | 5XBK | 3.22 |
| IPO5 | Importin-5 | 1096 | 6XTE | 2.27 |
| IPO9 | Importin-9 | 1041 | 6N1Z | 2.70 |
| IPO13 | Importin-13 | 963 | 2XWU | 2.80 |
| KPNA1 | Karyopherin subunit α-1 | 450 | 2JDQ | 2.20 |
| KPNA4 | Karyopherin subunit α-4 | 421 | 4UAE | 2.70 |
| KPNA6 | Karyopherin subunit α-6 | 479 | 4UAD | 2.42 |
| KPNB1 | Karyopherin subunit β-1 | 442 | 1F59 | 2.80 |
| TNPO1 | Transportin-1 | 890 | 2Z5J | 3.40 |
| TNPO3 | Transportin-3 | 923 | 4C0O | 2.56 |
| XPO1 | Exportin-1 | 1044 | 5DIS | 2.85 |
| XPO5 | Exportin-5 | 1204 | 3A6P | 2.92 |

**Fig. 1.**
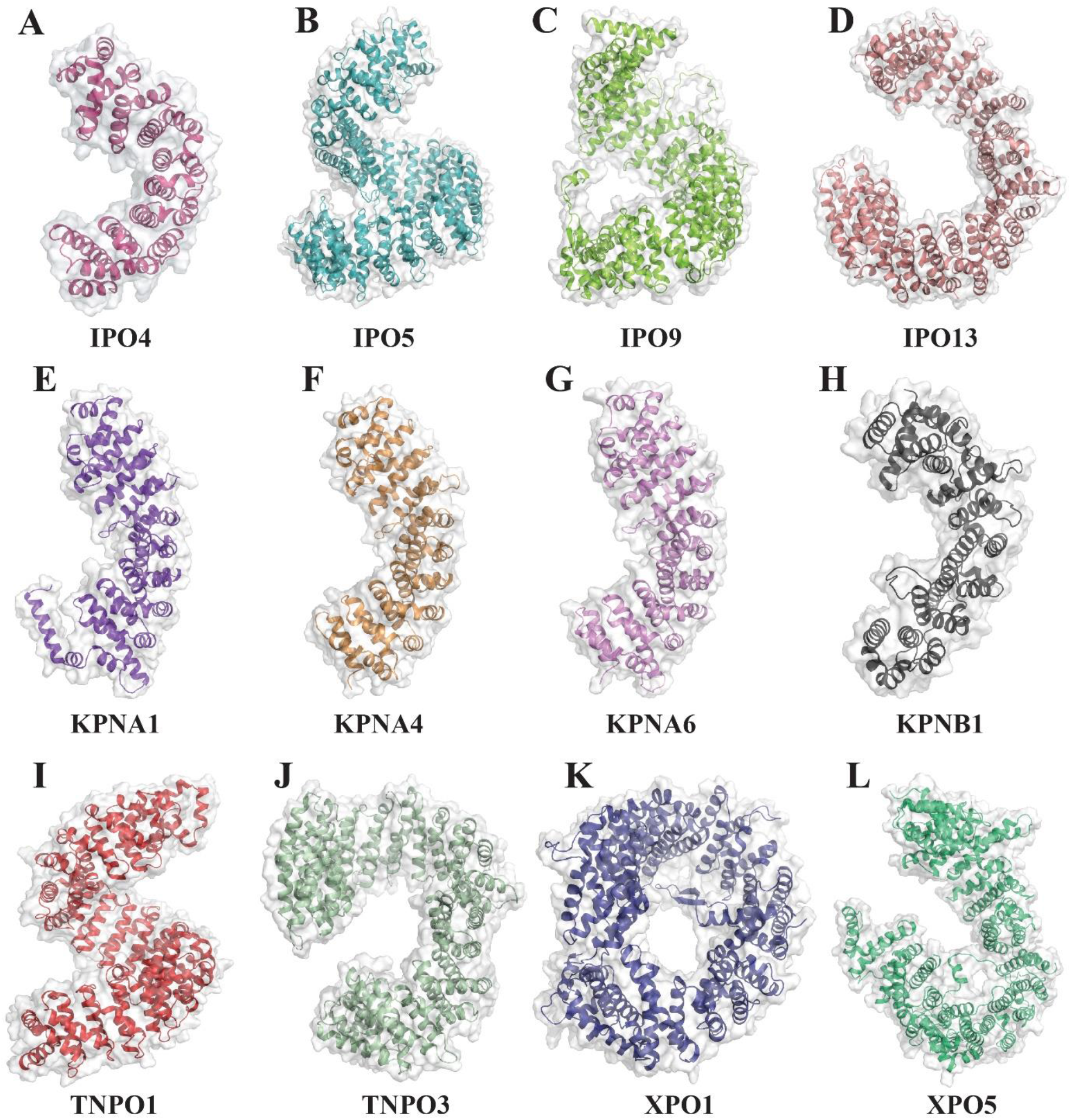
Cartoon and surface representation of 12 human nuclear transport proteins.

### Electrostatic potential analysis

To investigate the surface properties of the proteins, prepared proteins were uploaded to the eF-surf-PDBj (https://pdbj.org/eF-surf/top.do) web server and APBS Electrostatic plugin in PyMOL.

### Protein-protein docking

To investigate the protein-protein binding affinity and complex formation, molecular docking was performed at the widely used ClusPro v.2.0 (https://cluspro.org) web server [16]. ClusPro involves three important steps to dock protein structures, (a) rigid-body docking of Fast fourier transform (FFT) based generated billions of protein conformers using PIPER algorithm, (b) Root mean square deviation (RMSD) based clustering of 1,000 complexes and (c) refinement and energy minimization of complexes using CHARMM force field [17] to remove steric clashes between the interacting protein interfaces. To perform the molecular docking, two proteins were uploaded to the server as receptor and ligand based on sequence length.

Initially, each protein was docked with itself to form homodimeric complexes and, then docked with each other to get heterodimeric complexes. Hence, 12 homodimers and 66 heterodimers were obtained for further analysis. ClusPro gives best 30 complexes for each job after CHARMM energy minimization according to clustering probability and energy based (balanced, electrostatic-favored, hydrophobic-favored, and VDW+electrostatic-favored) parameters. In this study, best complexes were obtained by considering balanced energy parameter and higher clustering probability of complexes. The number of cluster members and the model cluster scores (cluster centre and lowest energy) are shown in Table S1. The cluster centre weighted score indicates the structure in the cluster that has the highest number of neighbour structures while the lowest energy score designates the structure which has the lowest energy in the cluster. This model score can be calculated using following equation;

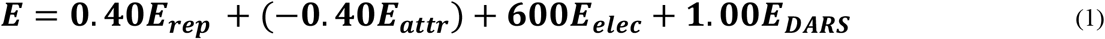

Where *E*_*rep*_ and *E*_*attr*_ indicate the repulsive and attractive terms of the van der Waals energy respectively whereas *E*_*elec*_ denotes electrostatic energy term. *E*_*DARS*_ is pairwise structure-based potential obtained by the decoys as the reference state approach [18] and represents the desolvation energy contribution. These docked complexes were further analysed for protein-protein binding affinity using PRODIGY [19] web tool available at https://wenmr.science.uu.nl/prodigy/. The binding energy evaluated at the PRODIGY server, was calculated using following equation [20],

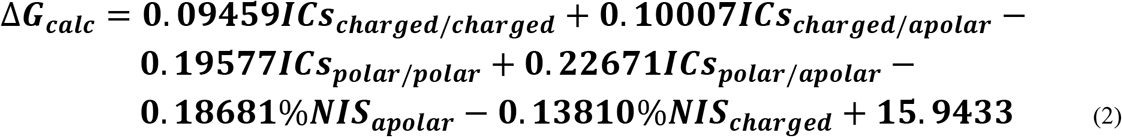

Where, *ICs* (Inter-residue contacts) and *NIS* (Non-interacting surface) terms indicate the contribution of various types of residues to the overall binding affinity.

### Protein-protein interface interactions analysis

To investigate the protein-protein interface interactions, PIMA [21] web server available at http://caps.ncbs.res.in/pima/. In the PIMA results, for the van der Waals energy, cut-off distance between protein atoms is 7 Å and the magnitude can be derived from the following equation,

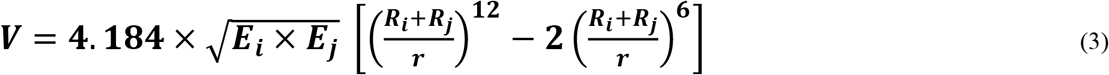

In the above equation, *R*_*i*_ and *R*_*j*_ are designated for the van der Waals radii of atom *i* and *j* whilst E represents the van der Waals well depth. The distance between two atoms is indicated by *r*. In the electrostatic energy (V) calculation, cut-off distance was set to 10 Å and energy can be calculated using following formula,

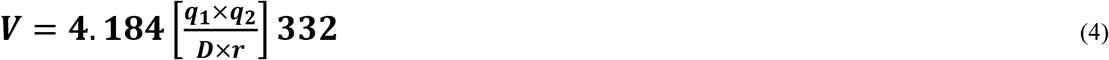

Where, *q*_*1*_ and *q*_*2*_ are partial charges of atom *i* and *j* while *r* is the distance between interacting two atoms.

Furthermore, PDBePISA available at https://www.ebi.ac.uk/pdbe/pisa/ and Arpeggio [22] available at http://biosig.unimelb.edu.au/arpeggioweb/ were employed to calculate various interactions and parameters between proteins in the complexes.

### Results and discussion Electrostatic potential analysis

Electrostatic interactions are considered to be an important driving force in various biomolecular mechanisms and protein-ligand or protein-protein bindings. Additionally, they also play crucial role in molecular recognition events [23]. Furthermore, for proper folding of nascent proteins and their stability, as well as in enzyme catalytic reactions, electrostatic forces play pivotal role [24]. Fig. 2 represents the electrostatic potential surface of each transport protein.

**Fig. 2.**
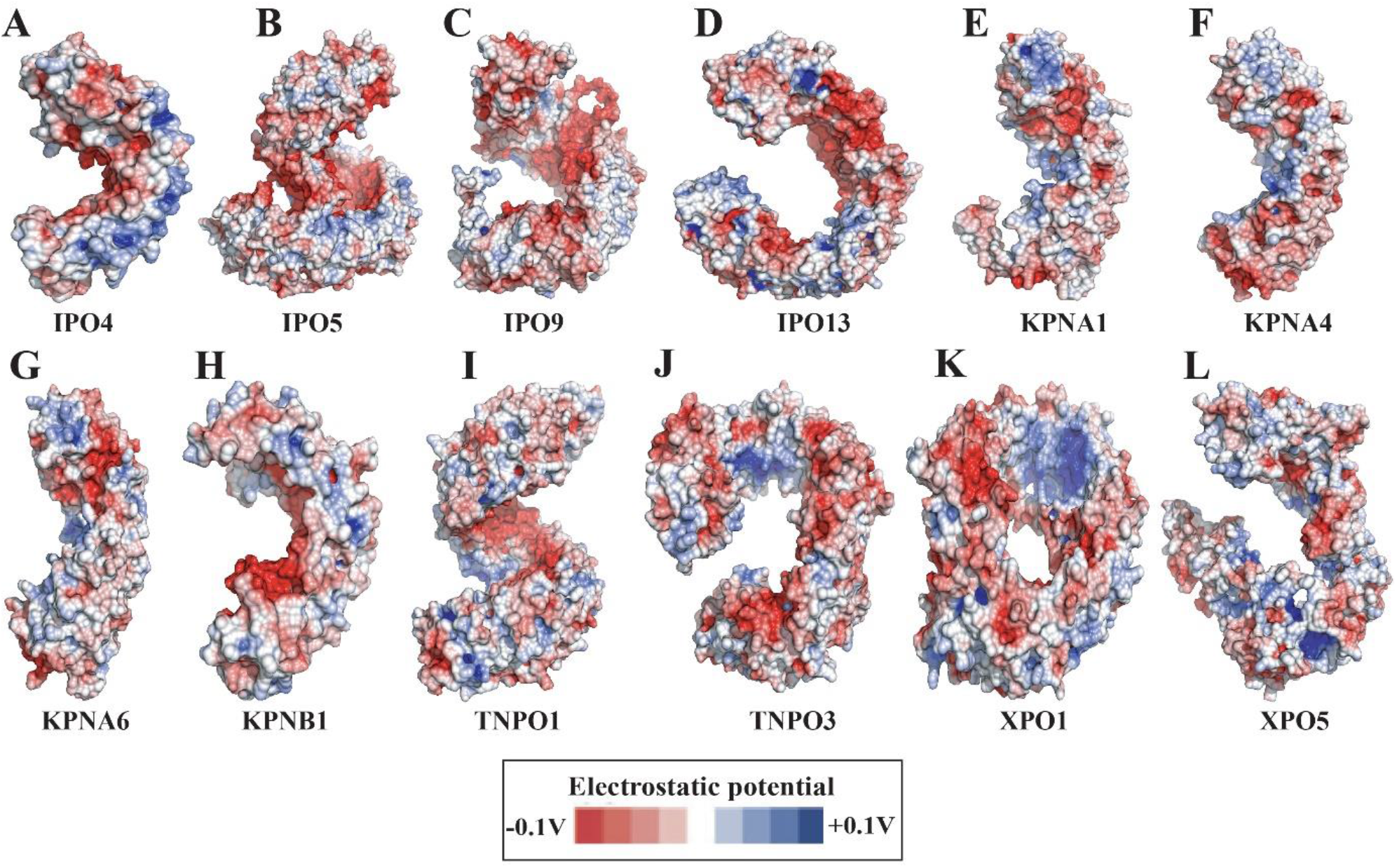
Electrostatic potential surface (ESP) representation of 12 human nuclear transport proteins.

In this study, to understand which regions of the protein have the specific charged or hydrophobic properties on the surface, Electrostatic potential (ESP) analysis was carried out. This analysis gives clue regarding the favourable protein sites where another protein binds. It can be seen from the Fig. 2 that in each protein, the internal surface of tandem repeats contains dominantly negative potential (red surface) whereas the outer surface has the significant amount of positively charged (blue surface) residues.

### Protein-protein docking

Protein-protein docking is used in various biomolecular applications such as, exploration of enzyme conformations [25], prediction of Interactomes [26], molecular recognition [27], protein dimerization [28], design of specific probes for protein targets [29], amyloid aggregation [30], predicting protein binding mechanisms [31], peptide designing against diseases [32] and vaccine designing [33]. In this study, binding energies and types of interactions of docked transport protein complexes (12 homodimers and 66 heterodimers) were evaluated using various computational tools. The calculated binding energies of complexes by PRODIGY web server are shown in Table 2 while the complex structures are illustrated in Fig. 3.

**Table 2.** Binding affinity (kcal/mol) of 78 human nuclear transport docked complexes calculated using PRODIGY web server.

| Complex number | Complex name | Binding energy (kcal/mol) | Complex number | Complex name | Binding energy (kcal/mol) |
| --- | --- | --- | --- | --- | --- |
| **IPO4** |  |  | 42 | IPO13-XPO5 | -13.7 |
| 1 | IPO4-IPO4 | -17.3 | **KPNA1** |  |  |
| 2 | IPO4-IPO5 | -10.4 | 43 | KPNA1-KPNA1 | -18.3 |
| 3 | IPO4-IPO9 | -16.4 | 44 | KPNA1-KPNA4 | -14.7 |
| 4 | IPO4-IPO13 | -13.8 | 45 | KPNA1-KPNA6 | -16.2 |
| 5 | IPO4-KPNA1 | -14.9 | 46 | KPNA1-KPNB1 | -21.9 |
| 6 | IPO4-KPNA4 | -12.5 | 47 | KPNA1-TNPO1 | -23.2 |
| 7 | IPO4-KPNA6 | -11.4 | 48 | KPNA1-TNPO3 | -20.0 |
| 8 | IPO4-KPNB1 | -12.5 | 49 | KPNA1-XPO1 | -16.3 |
| 9 | IPO4-TNPO1 | -16.0 | 50 | KPNA1-XPO5 | -14.0 |
| 10 | IPO4-TNPO3 | -14.4 | **KPNA4** |  |  |
| 11 | IPO4-XPO1 | -13.7 | 51 | KPNA4-KPNA4 | -11.7 |
| 12 | IPO4-XPO5 | -12.8 | 52 | KPNA4-KPNA6 | -12.0 |
| **IPO5** |  |  | 53 | KPNA4-KPNB1 | -17.7 |
| 13 | IPO5-IPO5 | -11.1 | 54 | KPNA4-TNPO1 | -15.3 |
| 14 | IPO5-IPO9 | -12.2 | 55 | KPNA4-TNPO3 | -15.9 |
| 15 | IPO5-IPO13 | -10.9 | 56 | KPNA4-XPO1 | -16.7 |
| 16 | IPO5-KPNA1 | -12.1 | 57 | KPNA4-XPO5 | -13.9 |
| 17 | IPO5-KPNA4 | -8.2 | **KPNA6** |  |  |
| 18 | IPO5-KPNA6 | -14.5 | 58 | KPNA6-KPNA6 | -12.0 |
| 19 | IPO5-KPNB1 | -7.7 | 59 | KPNA6-KPNB1 | -10.4 |
| 20 | IPO5-TNPO1 | -18.1 | 60 | KPNA6-TNPO1 | -12.7 |
| 21 | IPO5-TNPO3 | -15.1 | 61 | KPNA6-TNPO3 | -17.6 |
| 22 | IPO5-XPO1 | -13.2 | 62 | KPNA6-XPO1 | -15.7 |
| 23 | IPO5-XPO5 | -11.4 | 63 | KPNA6-XPO5 | -15.5 |
| **IPO9** |  |  | **KPNB1** |  |  |
| 24 | IPO9-IPO9 | -15.5 | 64 | KPNB1-KPNB1 | -6.6 |
| 25 | IPO9-IPO13 | -16.3 | 65 | KPNB1-TNPO1 | -13.8 |
| 26 | IPO9-KPNA1 | -15.1 | 66 | KPNB1-TNPO3 | -14.2 |
| 27 | IPO9-KPNA4 | -15.1 | 67 | KPNB1-XPO1 | -13.0 |
| 28 | IPO9-KPNA6 | -14.7 | 68 | KPNB1-XPO5 | -12.4 |
| 29 | IPO9-KPNB1 | -11.6 | **TNPO1** |  |  |
| 30 | IPO9-TNPO1 | -16.0 | 69 | TNPO1-TNPO1 | -19.4 |
| 31 | IPO9-TNPO3 | -12.7 | 70 | TNPO1-TNPO3 | -13.6 |
| 32 | IPO9-XPO1 | -14.2 | 71 | TNPO1-XPO1 | -18.2 |
| 33 | IPO9-XPO5 | -14.1 | 72 | TNPO1-XPO5 | -14.6 |
| **IPO13** |  |  | **TNPO3** |  |  |
| 34 | IPO13-IPO13 | -18.3 | 73 | TNPO3-TNPO3 | -14.7 |
| 35 | IPO13-KPNA1 | -16.9 | 74 | TNPO3-XPO1 | -11.7 |
| 36 | IPO13-KPNA4 | -15.3 | 75 | TNPO3-XPO5 | -11.8 |
| 37 | IPO13-KPNA6 | -13.6 | **XPO1** |  |  |
| 38 | IPO13-KPNB1 | -14.3 | 76 | XPO1-XPO1 | -11.2 |
| 39 | IPO13-TNPO1 | -16.9 | 77 | XPO1-XPO5 | -14.7 |
| 40 | IPO13-TNPO3 | -17.2 | **XPO5** |  |  |
| 41 | IPO13-XPO1 | -16.8 | 78 | XPO5-XPO5 | -10.9 |

**Fig. 3.**
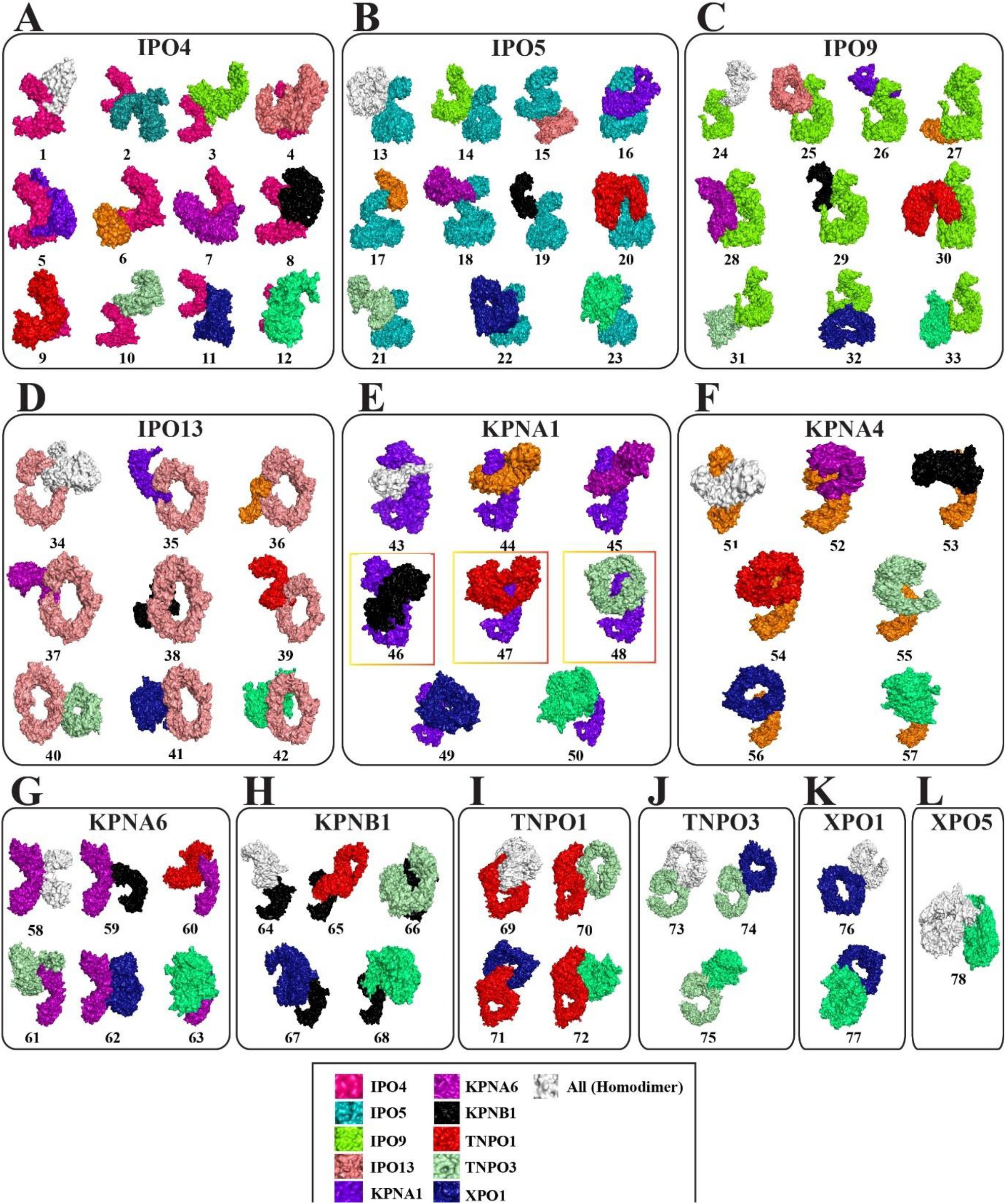
Docked 78 complexes (12 homodimers and 66 heterodimers) of 12 human nuclear transport proteins. The specific color is given to each protein in the complexes but in case of homodimer, the monomer is illustrated in white color. Top-three complexes (46: KPNA1-KPNB1, 47: KPNA1-TNPO1 and KPNA1-TNPO3) which have the highest binding affinities are shown inside the red-orange rectangular boxes.

Among 78 complexes, protein-protein binding affinity ranges from -6.0 kcal/mol to -23.5 kcal/mol (Fig. 4A). Additionally, it was noted that KPNA1, TNPO1 and TNPO3 have stronger affinity among 12 proteins to bind with their protein partners (Fig. 4B). Further, we screened complexes based on their binding affinity magnitudes and, we analysed top-three complexes, KPNA1-KPNB1, KPNA1-TNPO1 and KPNA1-TNPO3 (Fig. 5) for the interactions and energetics. Fig. 5A illustrates the 3D surface map with the projection of binding affinities of 78 complexes and energy distribution. And, Fig. 5B shows the heatmap of binding affinities of these complexes and reveals the strength of protein-protein binding.

**Fig. 4.**
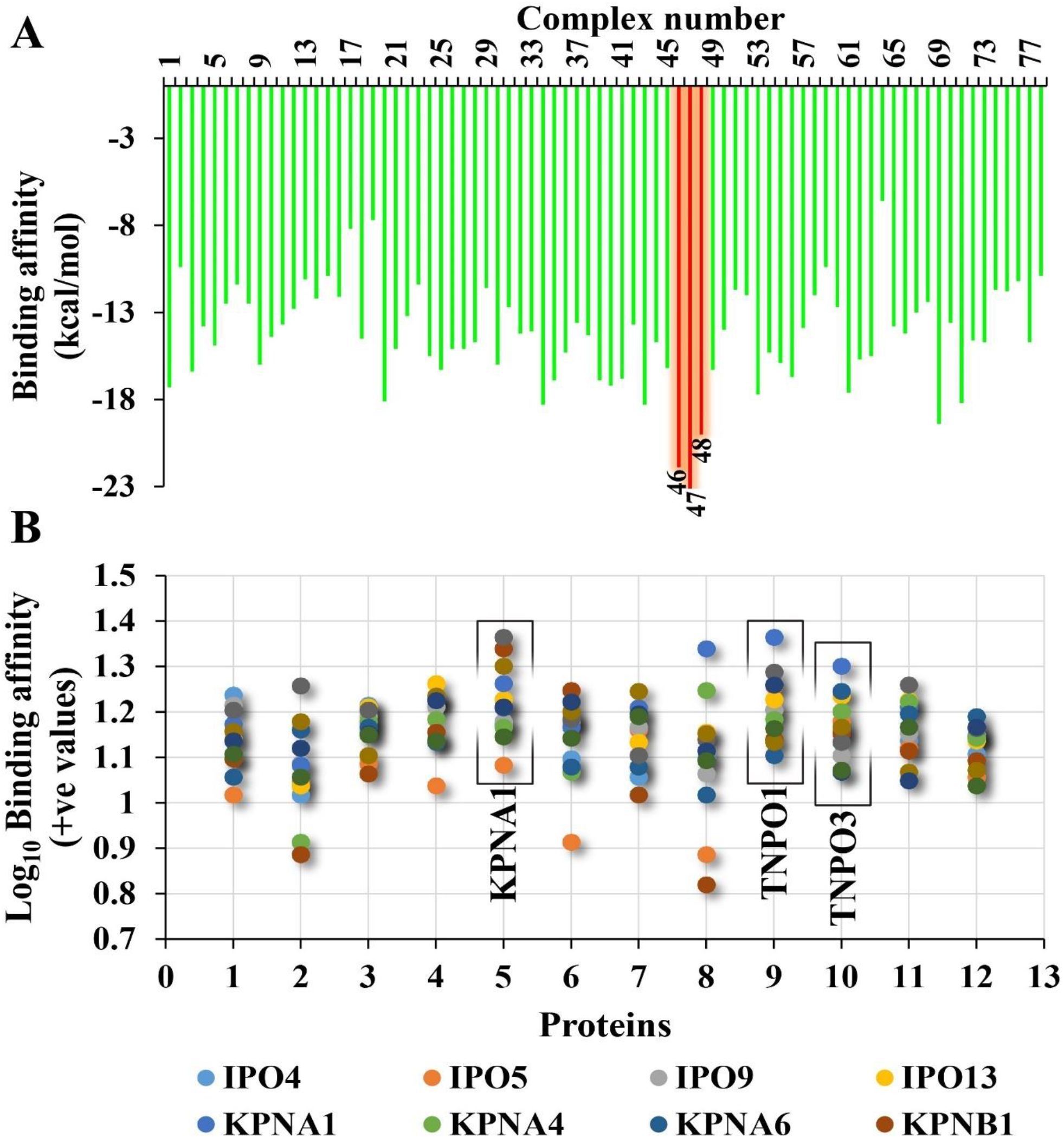
(A) Graphical representation of binding energy (kcal/mol) of 78 complexes calculated using PRODIGY web server. Read bars indicate the complexes which have higher binding affinity (46: KPNA1-KPNB1 (−21.9 kcal/mol), 47: KPNA1-TNPO1 (−23.2 kcal/mol) and 48: KPNA1-TNPO3 (−20.0 kcal/mol)). (B) Scatter plot represents the overall binding affinity of each protein with its protein partners along with itself. KPNA1, TNPO1 and TNPO3 which have higher binding tendency for their partners are indicated in black rectangular boxes.

**Fig. 5.**
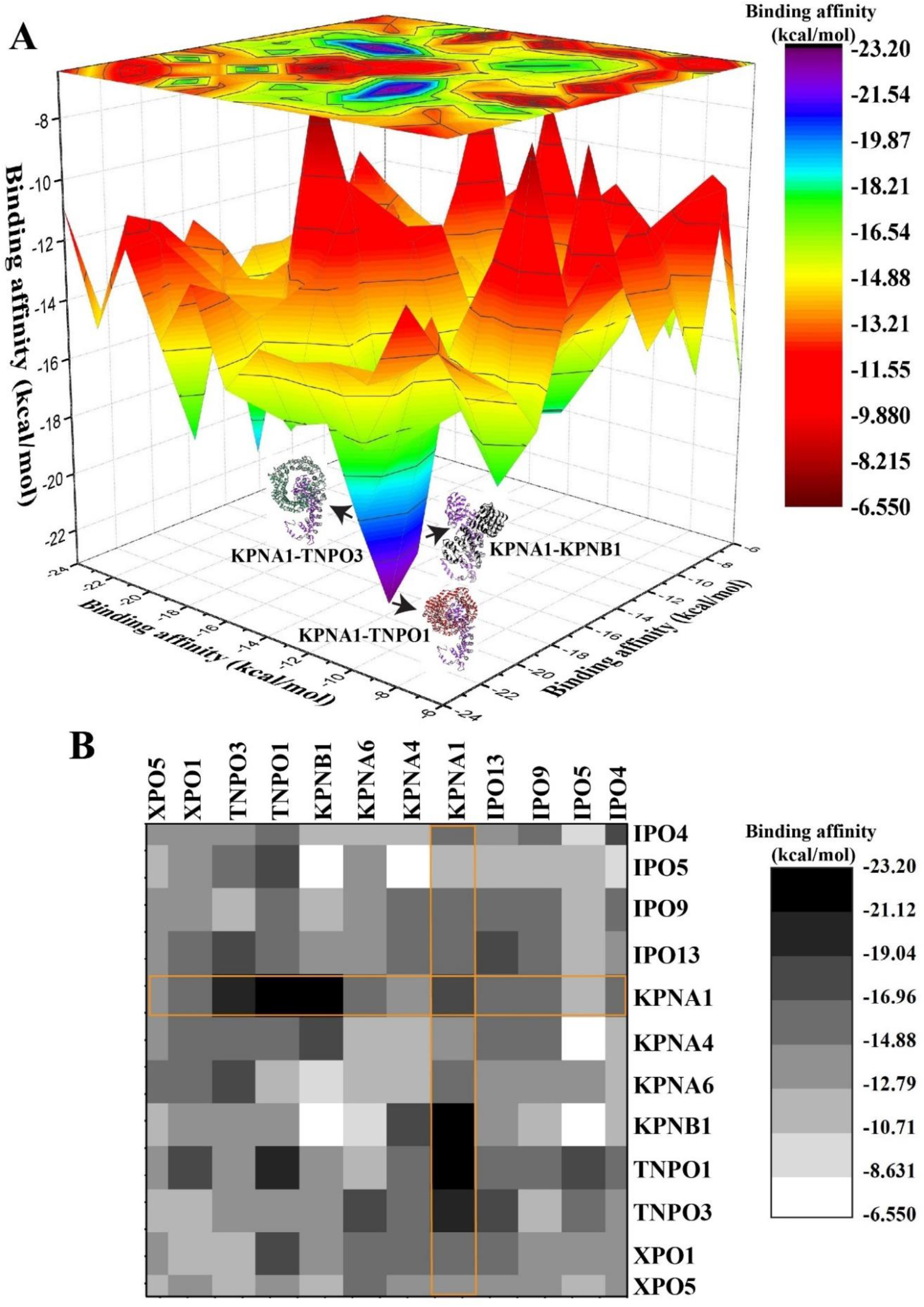
(A) 3D-surface map with a projection of binding affinities of 78 complexes. (B) Heatmap of 78 complexes. Yellow rectangular boxes represent the binding affinities of 12 KPNA1 complexes with other 11 proteins along with itself. Both plots are created using 12×12 matrix of binding affinities.

To further validate the results, we employed various tools to get insight into the protein-protein interactions profile of these three complexes (Table 3). Results indicated in the Table 3 support that KPNA1-TNPO1 complex has greater strength as compared to remaining two. The solvation energy is -7.8 kcal/mol that indicates the stability of this complex in the solvent due to greater number of hydrophobic residues are buried inside the complex. Furthermore, the total number of interacting residues of each protein, van der Waals (vdW), polar (hydrogen bonds & other), ionic (salt bridges) and hydrophobic (aromatic and alkyl) interactions are observed considerably higher than in comparison with other two complexes. And, protein-protein interface area is also significantly higher (2553 Å^2^) in KPNA1-TNPO1 as compared to 2479 Å^2^ and 2250 Å^2^ in KPNA1-KPNB1 and KPNA1-TNPO3 respectively. However, PIMA server predicts KPNA1-KPNB1 the most stable (−183.6 kcal/mol) complex in as compared to KPNA1-TNPO1 (−174.4 kcal/mol) and KPNA1-TNPO3 (−147.4 kcal/mol). Considering overall parameters, KPNA1-KPNB1 appears the second most stable complex among the 78 complexes. Analysis suggests that vdW interactions contribute enormously to the total stabilization energies of the complexes as compared to electrostatic interactions (Table 3).

**Table 3.** Protein-protein binding energies and interactions of top-three complexes, KPNA1-KPNB1, KPNA1-TNPO1 and KPNA1-TNPO3.

| Energy components & Contacts | Complexes |  |  |
| --- | --- | --- | --- |
|  | KPNA1-KPNB1 | KPNA1-TNPO1 | KPNA1-TNPO3 |
| Binding affinity (kcal/mol) | -21.9 | -23.2 | -20.0 |
| Total stabilization energy (kcal/mol) | -183.6 | -174.4 | -147.4 |
| Electrostatic energy (kcal/mol) | -82.8 | -69.9 | -55.9 |
| vdW energy (kcal/mol) | -100.7 | -104.5 | -91.4 |
| Solvation energy (kcal/mol) | 4.2 | -7.8 | -1.4 |
| N<sub>res</sub> (total) | 209 | 208 | 205 |
| Receptor-N<sub>Res</sub> | 73 | 78 | 75 |
| Ligand-N<sub>Res</sub> | 70 | 70 | 66 |
| Interface area (Å<sup>2</sup>) | 2479 | 2553 | 2250 |
| <sup>‡</sup>N<sub>vdW</sub> pairs | 5545 | 6223 | 5054 |
| *N<sub>vdW</sub> | 33 | 40 | 37 |
| N<sub>Polar</sub> | 135 | 159 | 123 |
| #N<sub>HB</sub> | 44 | 30 | 39 |
| *N<sub>HB</sub> | 81 | 98 | 69 |
| #N<sub>SB</sub> | 44 | 36 | 20 |
| <sup>‡</sup>N<sub>SB</sub> | 42 | 35 | 19 |
| N<sub>Ionic</sub> | 7 | 21 | 11 |
| N<sub>Aromatic</sub> | 9 | 10 | 9 |
| N<sub>Hydrophobic</sub> | 96 | 87 | 72 |
| N<sub>Carbonyl</sub> | 10 | 13 | 13 |

\*Arpeggio prediction #PDBePISA prediction <sup>‡</sup>PIMA prediction

Previous studies reported that KPNA1 interacts with KPNB1 [34-35]. But there are no structural level analyses have been delineated yet. Further, it is reported that TNPO1 does not require importin α also called karyopherin α (KPNA) subunits to transport it cargo proteins [36]. And, TNPO3 has not been reported yet to interact with any other transport proteins.

Protein-protein docking analysis shows that the KPNA1 harbours all three targets, KPNB1, TNPO1 and TNPO3 at the same site (Write a regions name) (Fig. 6-8). To understand the interactions profile of these three complexes, we carried out protein-protein interactions analysis. From the analysis, it was noted that acidic and basic residues dominantly participate to interact in the complexes. The total number of calculated hydrogen bonds and salt bridges between the protein in the complexes are given in Table S2. In the draft, we illustrated hydrogen bonds and salt bridges which are strong in nature due to observed at shorter distances.

**Fig. 6.**
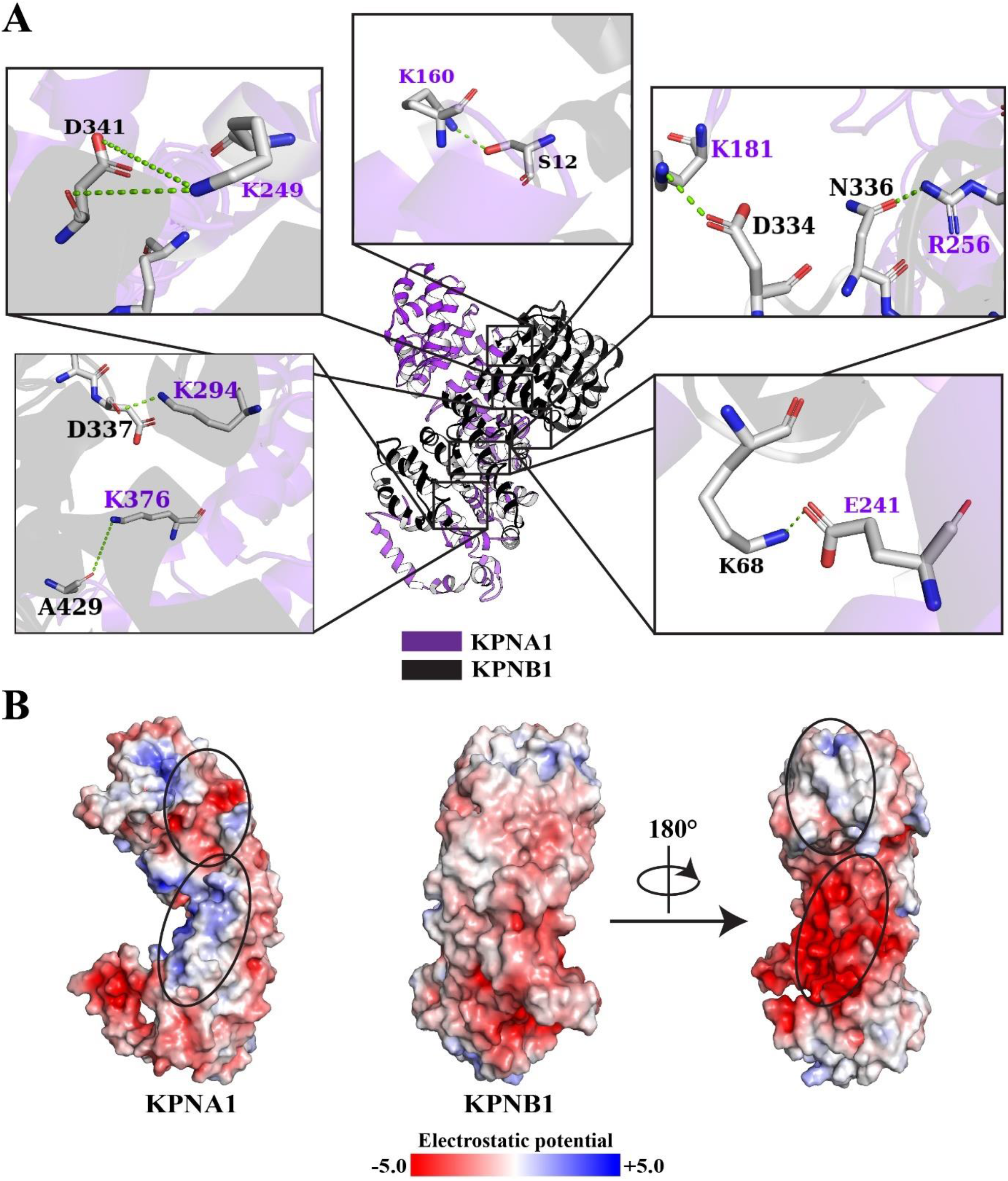
(A) Hydrogen bonds between KPNA1 and KPNB1 proteins in their complex (KPNA1-KPNB1). (B) ESP surface representation of KPNA1 and KPNB1. Protein-protein interface regions are shown in black circles.

In case of KPNA1-KPNB1 complex, D122, D205, D232, E51, E121, E241, K181, K249, K376, R163, R179, R238, R256, R333 and R335 residues from the KPNA1 and D334, D337, D338, D341, D426, E148, E437, E441, K62, K68, K269, R15 and R105 residues from the KPNB1 participate dominantly and form the hydrogen bonds and salt bridges to stabilize the complex. Fig. 6 represents the hydrogen bond interactions between the KPNA1 and KPNB1 in complex where only hydrogen bonds at the short distances are illustrated. The solvation energy is positive (4.2 kcal/mol) that indicates the slight destabilization of this complex into the solvent system. Its interface area is marginally less (2479 Å^2^) than the interface area (2553 Å^2^) of KPNA1-TNPO1 but, significantly higher than the KPNA1-TNPO3 which has 2250 Å^2^ (Table 3). Additionally, the hydrophobic interactions in KPNA1-KPNB1 were observed large in number as compared to other two complexes. Also, electrostatic contribution to the total stabilization energy of the complex was predicted greater (−82.8 kcal/mol) than the KPNA1-TNPO1 (−69.9 kcal/mol) and KPNA1-TNPO3 (−55.9 kcal/mol). And, from the ESP analysis, it can be seen from the Fig. 6B that KPNA1 has negatively charged surface at the upper neck and positively charged residues at the middle regions. Hence, KPNB1 has complementary potential surface to bind with KPNA1.

Next, KPNA1-TNPO1 complex was predicted the most stable complex among the all complexes. Most of the determined parameters predict this complex most stable except some parameters like total stabilization energy and hydrogen bonds calculated using PIMA and PDBePISA respectively (Table 3). Interactions analysis revealed that D211, D247, E7, E125, R45, R68, R101, R163 and R179 residues of KPNA1 and E7, E285, E510, E682, K290, K319, R246, R295 and R391 residues of TNPO1 largely contribute to the total interactions which results into stability of this complex among all the complexes. Additionally, this complex has considerably large number of hydrogen bonds, van der Waals and ionic interactions and, slightly higher number of carbonyl and aromatic interactions as compared to KPNA1-KPNB1 and KPNA1-TNPO3 complexes (Table 3). The stronger interactions at the short distances between the KPNA1 and TNPO1 are illustrated in Fig. 7. Additionally, ESP analysis suggests that the middle region of the TNPO1 which is negatively charged and binds at the neck region of KPNA1 which has positive potential (Fig. 7B).

**Fig. 7.**
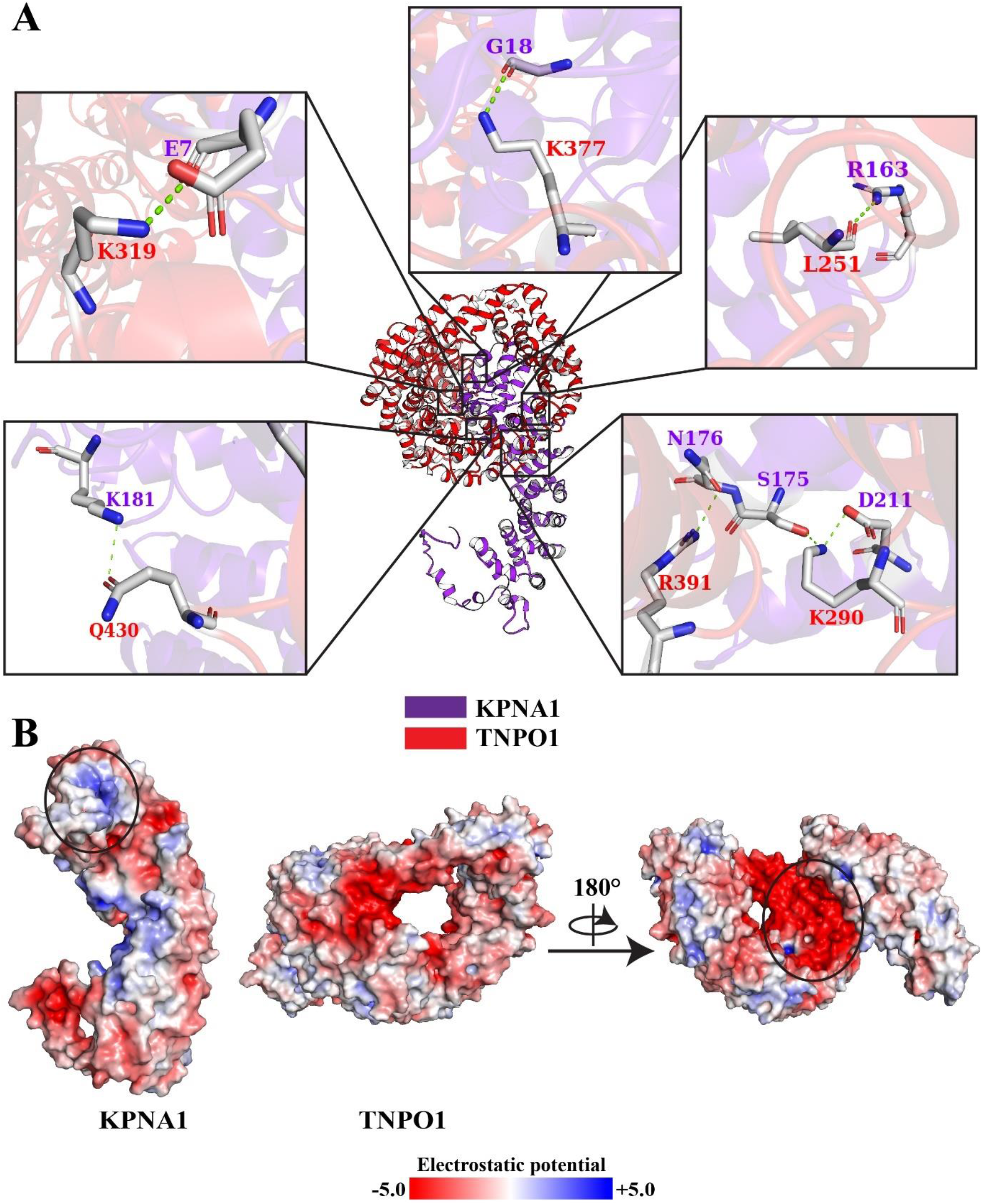
(A) Hydrogen bonds between KPNA1 and TNPO1 proteins in their complex (KPNA1-TNPO1). (B) ESP surface representation of KPNA1 and TNPO1. Protein-protein interface regions are shown in black circles.

Finally, considering the third complex, KPNA1-TNPO3 is the third most stable complex. Comparing the energetics and protein-protein interactions in this complex to the KPNA1-KPNB1 and KPNA1-TNPO1 indicates that this complex has significant amount of less binding affinity and interactions. However, solvation energy (−1.4 kcal/mol) indicates slightly stable into the solution (Table 3). Fig. 8 illustrates the proximal and strong interactions between the KPNA1 and TNPO3. Further, from the detailed interactions analysis, it was noted that residues, D137, D211, D232, E51, E114 and N195 in KPNA1 and E28, E257, R165, R166, R348, R789,

**Fig. 8.**
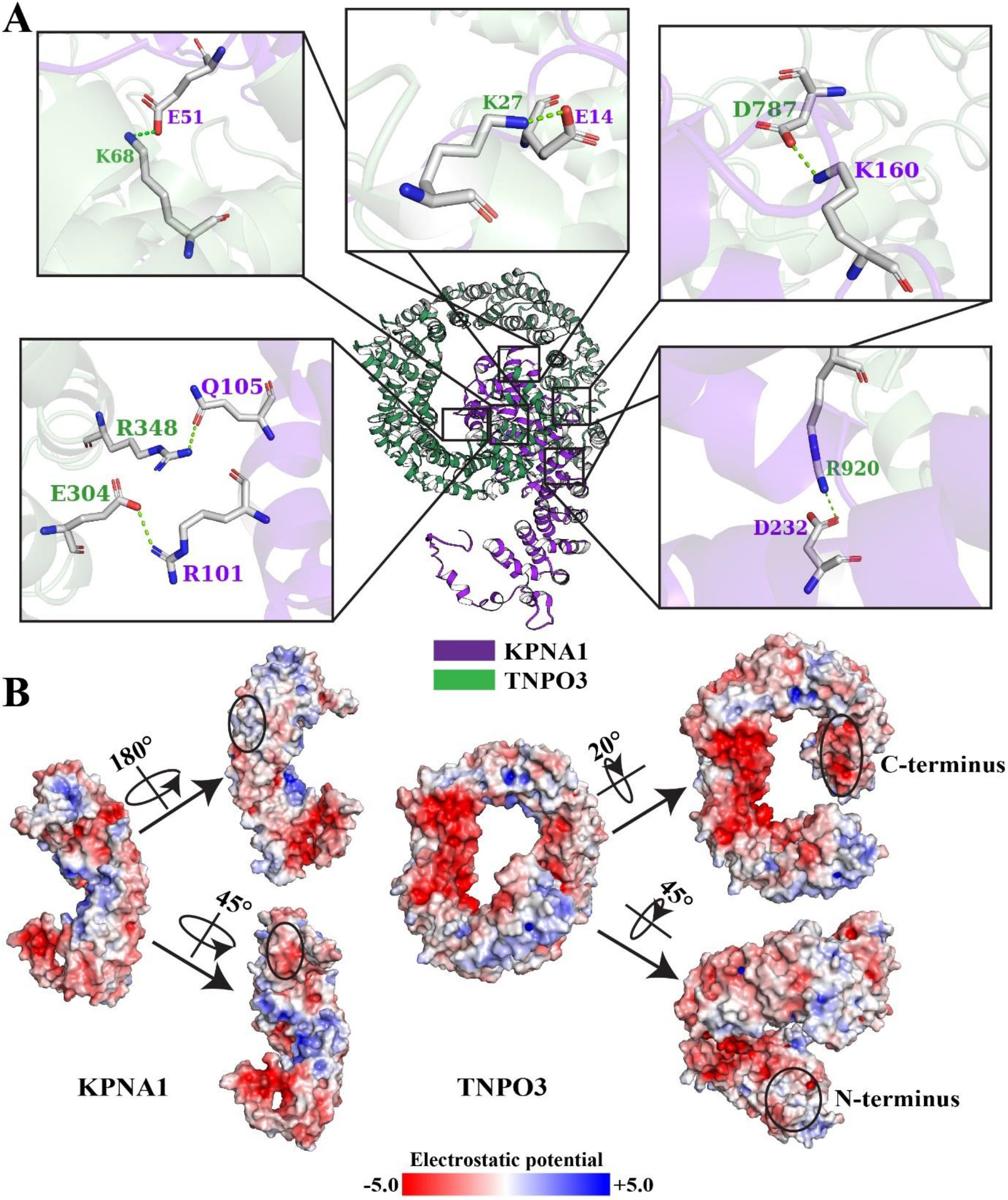
Hydrogen bonds between KPNA1 and TNPO3 proteins in their complex (KPNA1-TNPO3). (B) ESP surface representation of KPNA1 and TNPO3. Protein-protein interface regions are shown in black circles.

R920 and R923 residues in TNPO3 essentially contribute to the stability of this complex (Table S2). In this complex, TNPO3 binds to the KPNA1 at the neck regions like TNPO1 through its N-and C-terminal regions. The inner region of the N-terminus is slightly positively charged whereas C-terminus is negatively charged. Thus, the outer region of the KPNA1 at the top left is negatively charged where N-terminus of the TNPO3 is located while the middle back outer region is positively charged where C-terminus of the TNPO3 is positioned (Fig. 8B).

## Conclusion

The primary objective of this study was to predict whether the transport proteins have tendency to interact and bind with each other. As it is a fact that in the thronging and dynamic cellular environment, proteins have specific and nonspecific or transient interactions between the DNA-protein, RNA-protein, protein-protein, protein-lipid and protein-small molecule. Thus, to determine the binding affinities and the nonspecific interactions between 12 human nuclear transport proteins, protein-protein docking study was performed using ClusPro web server. Each protein was docked with itself and remaining 11 proteins hence, the total 78 complexes (12 homodimers & 66 heterodimers) were generated. Further, multiple web tools such as PRODIGY, PIMA, PDBePISA and Arpeggio were employed to analyse complexes for their interactions and energetics. Results showed that three proteins, KPNA1, TNPO1 and TNPO3 have the greater potential to bind with other proteins. And, three complexes (KPNA1-KPNB1, KPNA1-TNPO1 & KPNA1-TNPO3) of KPNA1 were observed to have higher stabilities among 78 complexes. This study is primary investigation thus, further biophysical analyses are necessary to confirm the above findings.

## Acknowledgments

Author is thankful to his chemistry department for providing computational and infrastructure facilities.

